# Plasticity Dynamics Improves Reservoir Performance on Mixed-Timescale Memory Tasks

**DOI:** 10.64898/2026.09.24.754073

**Authors:** Marcelo P. Becker, Michael Fauth

## Abstract

Mixed-timescale signals are ubiquitous in natural stimuli and real-world applications, yet they remain challenging to process with recurrent neural networks. Reservoir computing offers an efficient framework for temporal processing, but extending its temporal range typically requires tuning neuronal timescales, which can compromise the representation of faster dynamics. Here, we propose an alternative approach in which synaptic plasticity provides an additional, slower timescale for computation. We introduce Hebbian plasticity into recurrent networks and show that its dynamics enable the reservoir to simultaneously process fast and slow components of mixed-timescale inputs: neuronal dynamics encode fast fluctuations, while synaptic dynamics support slower components. Remarkably, the resulting computational benefit extends over more than an order of magnitude beyond the intrinsic timescale of the plasticity. We show that this extended processing range arises from the dependence of the plasticity rule on postsynaptic activity, which effectively injects a filtered memory of past states into the network current. These results identify synaptic plasticity as a mechanism for extending the temporal processing capabilities of recurrent networks without altering neuronal timescales, and also suggest a potential computational role for plastic synapses in the processing of slowly varying stimuli in biological neuronal networks.

## 1 Introduction

Timeseries processing and forecasting is an important and active field of research in machine learning (Kim et al., 2024) and in neuroscience (Mauk and Buonomano, 2004). It can serve as a basis for decision-making problems from an engineering point of view, extracting information that is relevant for a particular task, or from the view of how the brain processes everyday stimuli. One approach to this problem that touches both interfaces is the reservoir computing one. It is a biologically inspired machine learning framework that utilizes recurrent neural networks to perform computations over a given input (Jaeger, 2001; Maass et al., 2002; Lukoševičius and Jaeger, 2009). In its echo state network formulation (ESN), it consists of a 3-layer structure of rate neurons: an input layer, a randomly connected recurrent layer, and an output layer that is trained to perform a task. The major advantage of this framework is that a costly training of the network’s recurrent weights is unnecessary. Instead, this approach uses the large repertoire of computations already present in the randomly connected recurrent network and only trains the output layer via linear regression, which is computationally cheap. Most of the standard benchmarks historically used, however, are designed with single-timescale tasks and uncorrelated inputs in mind (as in the case of delay and parity tasks), or involve mixing timescales of the immediate past (NARMA-10, chaotic systems prediction) (Jaeger, 2001; Cramer et al., 2020; Ozturk et al., 2007; Obst et al., 2010). Many of the real-world signals, however, contain information simultaneously at fast and slow timescales, requiring the model to retain recent inputs while integrating information over much longer periods, as is the case of language and music processing, and motor planning. When dealing with longer timescales, it is a common strategy to extend the reservoir’s dynamical range by introducing a leak parameter corresponding to the neurons’ intrinsic timescale (Jaeger et al., 2007). This, however, makes the network less effective in performing shorter-timescale tasks. Mixing shorter and longer timescales, then, can be challenging in the standard formulation of ESN’s.

Several approaches to solve this problem have been proposed in recent years. The most straightforward solution is to include changes in the neuronal leak rate. This can come in the form of heterogeneous leak rates which need to be tuned for the task (Tanaka et al., 2022), adaptable leak rates that use the properties of the dynamics to adapt (Dasgupta et al., 2013), or multiple reservoirs with tuned leak rates linked together in a hierarchical (Manneschi et al., 2021) or deep structure (Gallicchio et al., 2018). Another solution consists of introducing delays in the updating of the recurrent units, tuning the delays appropriately (Lun et al., 2016; Ma et al., 2021). Each of those solutions has its uses, performing better in some signal processing tasks or sometimes in the classification of multi-timescale stimuli, requiring evaluation of their use cases.

Here we follow an alternative approach which is inspired by biological neural circuits, which naturally process information across multiple temporal scales (for example, in working memory tasks (Brennan and Proekt, 2023; Wolff et al., 2017), and planning and decision making (Sutton, 1995; Tang et al., 2021)) despite individual neurons exhibiting relatively short intrinsic time constants of tens to hundreds of milliseconds Murray et al. (2014). Here, physical constraints prevent the neurons from increasing their leak rate to operate on slower timescales. However, because synaptic efficacy evolves more slowly than membrane potentials, synapses can store information beyond the intrinsic integration time of individual neurons (O’Donnell, 2023) or populations. We here propose that these *slower timescales of the synaptic dynamics can be integrated into reservoir computing to extend the dynamical repertoire* of the recurrent network.

We demonstrate that reservoirs employing this mechanism can solve mixed-timescale processing problems without modifying the neurons’ leak rates. For this, we equipped our echo-state network with a form of Hebbian plasticity with an explicit decay time constant and demonstrate that this improves computations on mixed-timescale delay and correlated NARMA tasks. We prove that this improvement indeed comes from the dynamics of the weights and not just an adjustment of the weights to a new static value. This comes with only a small computational cost that scales quadratically with the network size (or even linearly for simpler plasticity rules). We further provide a theoretical explanation for this improvement by showing that plastic synapses introduce a filtered memory term into the current while still exhibiting the leak neuronal timescale.

## 2 Results

### 2.1 Model and tasks overview

We investigate the computational capabilities of an echo-state network with synaptic plasticity using complex regression tasks on mixed-timescale inputs. To this end, we constructed a network with three distinct layers: an input layer consisting of multiple Ornstein-Uhlenbeck (OU) processes with different correlation timescales, a recurrent plastic layer, and a single-neuron readout. The recurrent layer is composed of time-discrete neurons with a sigmoidal (Fermi) non-linearity and no memory (100% leak every timestep). This corresponds to setting the time in units of the neuronal timescale, and considering the dynamics sufficiently smooth in this range. Recurrent connections are sparse and have a fixed Gaussian component with a given spectral radius, and a plastic Hebbian component (Fig. 1) characterized by a timescale *p* and an amplitude *k* similar to the ones studied by Dong and Hopfield (1992) and more recently by Clark and Abbott (2024). Positive values of *k* correspond to standard Hebbian plasticity, with pairwise correlations reinforcing connections, while negative *k* results in anti-Hebbian plasticity. For *k* null, we fall back to the non-plastic basic network, serving as a control case.

**Figure 1.**
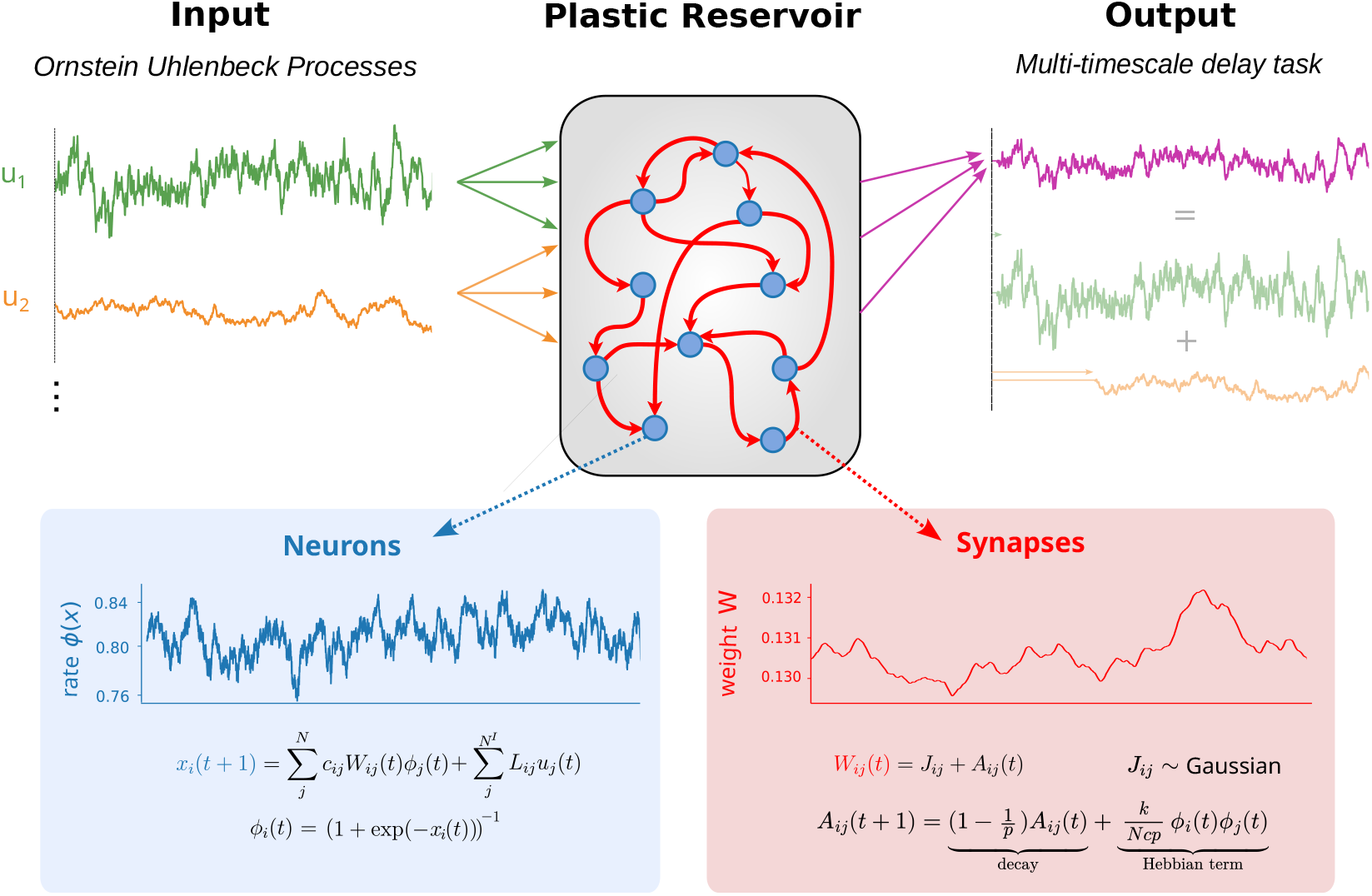
Diagram of the reservoir setup. We used an echo-state network composed of Fermi neurons with synapses comprised of a static component, drawn from a Gaussian distribution, and a plastic component. Inputs are drawn from Ornstein-Uhlenbeck processes with different correlation times. The main tasks used throughout this work where composed of reproducing averaged delayed versions of the inputs. The output layer is trained by performing a Tikhonov/Ridge regression over the desired target.

To evaluate the network’s ability to perform the memory computations, we used two different tasks: a delay task and a correlated NARMA-100 task. The delay task consists of calculating the average of delayed versions of the input, while the NARMA task incorporates a nonlinear product of previous timesteps of itself (Fig. 1). Training of the output is done by performing a Tikhonov/Ridge regression (see Methods).

### 2.2 Plasticity effects on mixed-timescales memory tasks

We first study the influence of plasticity using a delay task with two OU inputs – one fast and one slow (*τ*_1_ = 20 and *τ*_2_ = 400). As the target output, we use the sum of delayed versions of these signals, where the delays *d* were chosen to correspond to the timescale of the respective inputs (*d*_1_ = 20 and *d*_2_ = 400). Hence, the network should remember the more recent past of the fast input and a more distant one for the slow. We evaluated the NRMSE while varying the spectral radius of the static Gaussian weight matrix and the plasticity strength *k*.

We observe that plasticity does indeed influence the performance of the reservoir network on the delay task. When plasticity is active, that is, for *k* ≠ 0, the average NRMSE is substantially smaller than for the non-plastic case (*k* = 0) (Fig. 2A and D). The most notable performance gain, however, is observed at positive values of *k*, while for negative values the gain is somewhat smaller. As visible in the sampled output trajectories (Fig 2B), this improvement comes mostly from the Hebbian network being better able to appropriately tell when large fluctuations of the target occur. This view is further supported by the autocorrelation of the difference between output and target (Fig. 2C), which decreases faster for the *k* = 2 case (red curves), indicating that the majority of the error is concentrated in the faster modes of the target. On the other hand, *k* = −2 (green curves) cuts part of the longer lag components of the error when compared to the non-plastic case.

**Figure 2.**
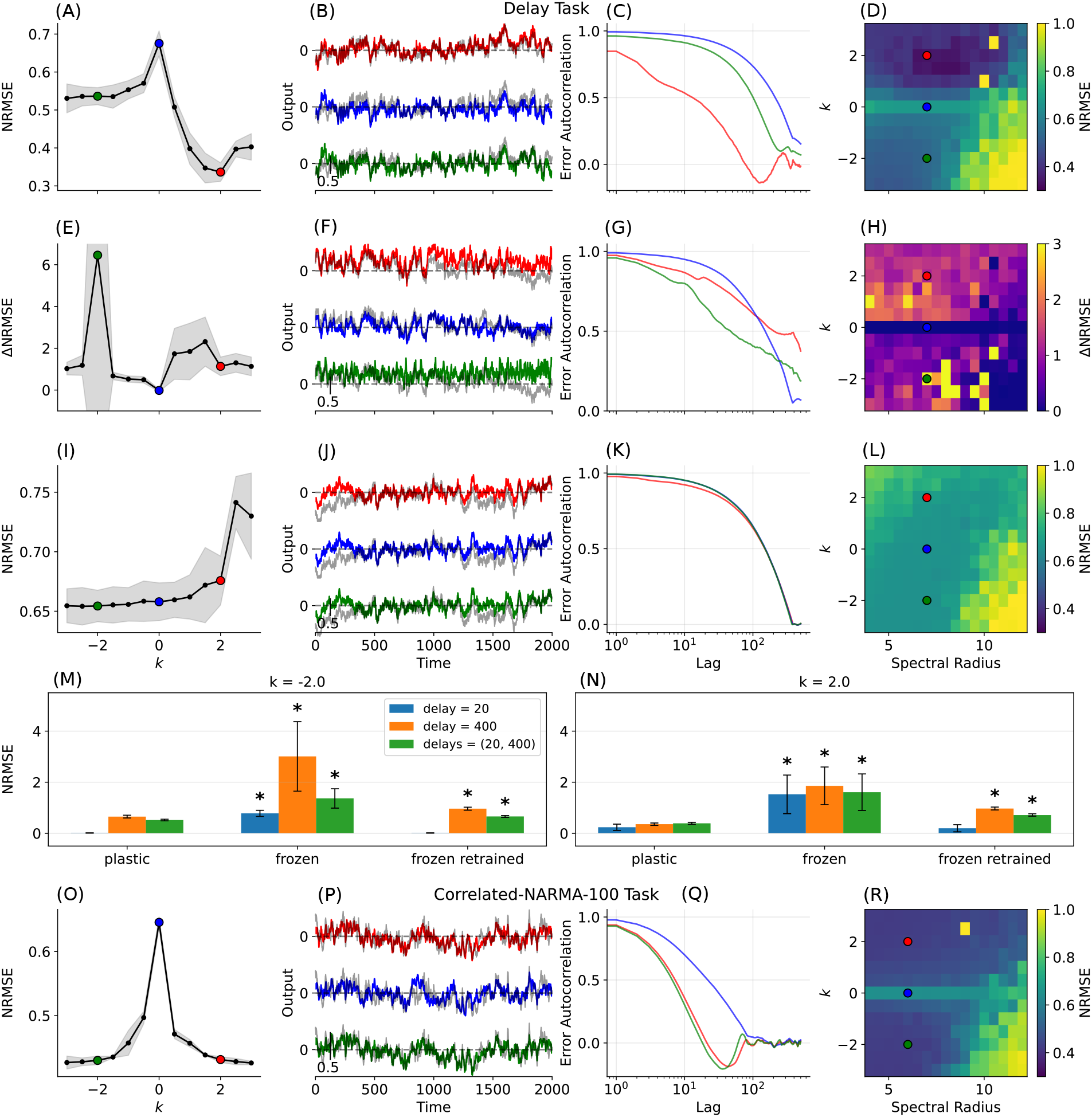
Plasticity allows reservoirs to solve mixed-timescales tasks. (A)-(D) Performance of networks on the delay task for different values of plasticity strength *k* and spectral radius *ρ*(*c*_*ij*_*J*_*ij*_). (A) The average of normalized root mean squared error (NRMSE) mean with *ρ*(*c*_*ij*_*J*_*ij*_) = 7. (B) Output traces of an interval of 2000 timesteps after training against the target output. Colors correspond to parameter values in (D). (C) Autocorrelation of the difference between the output trace and target in the testing phase. Autocorrelations were averaged over all realizations of the network and task. (D) NRMSE heatmap for *k* an *ρ*(*c*_*ij*_*J*_*ij*_). Values were capped at 1 for readability. (E)-(H) Similar to the above, for networks after freezing the plastic weights. ΔNRMSE gives the difference between the NRMSE after and before freezing the weights. (I)-(L) Figures for the frozen-weights network after retraining the output layer. Input consists of two OU processes with correlation times *τ* = 20 and *τ* = 400 and output calculated using delays *d*_1_ = 20 and *d*_2_ = 400 (1 to 1 ratio with inputs). Statistics were calculated over 5 realizations of each parameter combination. (M)-(N) Mean NRMSE for delay tasks with a single fast or slow timescale and the mixed one for the plastic, frozen, and frozen-retrained phases. Statistics are taken over 20 realizations. Stars indicate that the corresponding distribution is significantly higher than the corresponding plastic one (Mann-Whitney U test, *p <* 0.01). (O)-(R) Similar to (A)-(D) but for the correlated NARMA-100 task. Inputs consisted of 4 OU processes with correlation timescales *τ*_1_ = 10, *τ*_2_ = 30, *τ*_3_ = 70, *τ*_4_ = 100. Statistics taken over 3 realizations.

The dependence of the effect on the spectral radius is less sensitive. Setting up the network in a stable regime is mostly sufficient to get the positive temporal effects of plasticity. There is, however, a small region in the parameter space where the network behaves optimally for this specific task configuration. We note that for other choices of input and output timescales that we tested, this minimum was not present (data not shown). The location and even the existence of this region is then dependent on the properties of the input and task. For stronger values of the spectral radius, we observe the complete degradation of the network’s ability to perform the task. This appears to happen when we cross a chaotic boundary, where small perturbations grow and make linear readability not possible. This is mostly visible in the anti-Hebbian case.

These results indicate that plasticity does indeed improve performance on delay tasks. Yet, it still needs to be clarified whether the performance improvements actually come from the intrinsic dynamics of the plastic weights or simply arise from a convergence to an optimal set of weights. To disentangle these two possible explanations, we carry on the simulation after the testing phase but with the recurrent weights frozen. In this way, if the network performance comes from an optimal weight fixed-point, we expect the overall error not to change significantly. This is not what we observe, though (Fig. 2E-H): freezing the weights degrades the performance of the plastic networks to the point that the network performs worse than predicting the mean (ΔNRMSEs goes beyond one). In regions where the error was already large, no change is observed. The error autocorrelation in this case shows the progressive degradation effect of freezing the weights, with the autocorrelation exhibiting a longer tail than the *k* = 0 case, spreading the error over longer timescales (Fig. 2(G).

We also retrained the outputs after freezing the weights to see if any further improvements could be achieved in this setting. After this retraining, the network performance is comparable to that from the original non-plastic network (Fig. 2I-L). Thus, it is clear that the linear readout is indeed using the plasticity dynamics to perform its memory task. This means that the plasticity timescales show their characteristics in the dynamics of the neurons in a readable manner. It can therefore be assumed that our network has access to both the neuronal and the plasticity timescales, which makes it especially suitable for mixed-timescale computations such as the above-described multiple delay.

In particular, we hypothesized that each of the delays in our task is solved by one of the dynamic timescales of the reservoir. To test this, we compare the error of the three above-described stages for tasks involving only the fast timescale (20), only the slow timescale (400), and both together (20 and 400). Statistical significance was calculated using a Mann-Whitney U test (*p <* 0.01) to see if the error in the frozen and frozen-retrained protocol was higher than the plastic one. For *k* = −2, the plastic network can handle the fast task better than the slow one. It exhibits a smaller degradation for it when freezing the weights and regains its performance after retraining (Fig. 2M). This indicates that most of the work in the fast task is done using the neuronal dynamics, not the plasticity. This effect is clearer in the *k* = −2 case. The plastic network can solve all three cases with a small NRMSE. After freezing the weights, all tasks show major degradation, but only for the fast task does the performance recover completely after retraining (Fig. 2N). Thus, the network is using the neuronal dynamics to tackle the fast part of the task and the plasticity dynamics to handle the slow part.

Finally, we tested whether the effect of plasticity is conserved with the introduction of nonlinear dependencies between timescales. For this, we evaluated NRMSE and error autocovariance for a NARMA-100 task. We indeed observed similar improvements for plastic networks (Fig. 2O-R). However, in contrast to the multiple delay task, error levels for both Hebbian and anti-Hebbian networks in the stable regime and the error autocovariance are comparable.

### 2.3 Influence of plasticity timescales on the delay task

We have shown that synapse dynamics introduced by plasticity are able to extend the network timescales in a way that is readable by an output layer performing a mixed timescale task. This leads us to ask: how does the plasticity timescale *p* influence the processing of the inputs with different timescales (correlation times of the OU-processes described above)? To answer that, we again used the multiple delay task, but now varying the plasticity timescale while fixing the plasticity strength at *k* = 2. Within a large range of the spectral radii, we find a broad optimal region for the synaptic timescale (Fig. 3A-D). Too small values of *p* bring the plasticity timescale closer to the neuronal timescale and render the network unable to solve the task reliably, with errors close to the non-plastic case. For larger values of *p*, the performance slowly degrades. These effects can be seen directly in the output traces (Fig. 3B) and the error autocorrelation (Fig. 3C). For smaller *p*, the network does not properly account for the larger, slower fluctuations of the target signal, causing the trace to trail off the target. For intermediary values, this problem is largely corrected, with the error autocorrelation exhibiting the fastest decay. However, for even larger values of *p*, the output trajectory is overly smoothed out, suppressing higher-frequency fluctuation components and generating a higher error autocorrelation in the lower lag range. This indicates that the plasticity timescale should be matched to input and task properties.

**Figure 3.**
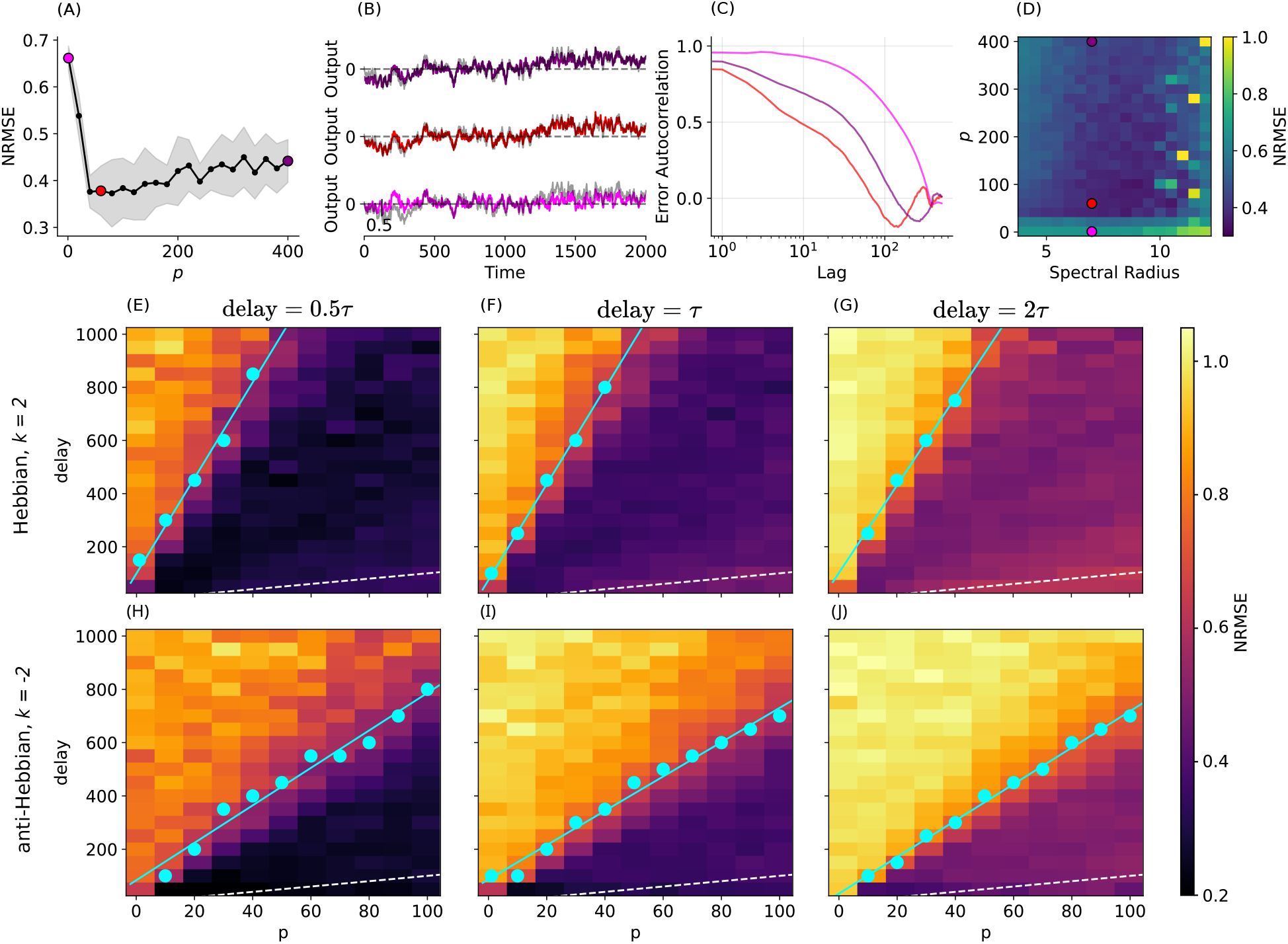
Possible delays are orders of magnitude larger than the plasticity timescale. (A) NRMSE for different plasticity timescales *p* and spectral radius *ρ*(*c*_*ij*_*J*_*ij*_) = 7, averaged over 5 realizations, with error bars calculated as standard deviations. (B) Output traces for networks with Hebbian plasticity (*k* = 2) for three values of the synaptic timescale *p*, with the target output in grey. Colors correspond to parameter values in (D). (B) NRMSE averaged over 5 realizations of the network and input. Colored dots correspond to the curves in (A). (C) Autocorrelation of the difference between the output trace and the target. Autocorrelations are averaged over all the realizations. (D) NRMSE heatmap for *p* against the spectral radius. Values are capped at 1. (E)-(G) NRMSE heatmaps averaged over 5 repetitions for different plasticity timescales *p* and task delays with *ρ*(*c*_*ij*_*J*_*ij*_) = 6. We also scale the input correlation with the delay, using the ratios 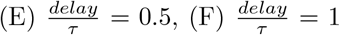 and 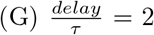. (H)-(J) same as above, but with anti-Hebbian plasticity (*k* = −2). Inputs consisted of a single timeseries with the referred correlation, and the target was a delayed version of this input. Dots correspond to the point where the average NRMSE crosses the half-point for that value of *p*. A line was fitted to the points giving the slope and intercept pairs (D) (17.4, 119.3), (E) (17.9, 79.2), (F) (16.5, 100.0), (G) (7.0, 83.3), (H) (6.4, 86.6), and (I) (6.8, 33.3). The traced line is the identity line.

To better understand the relationship of these three timescales, we restrict the inputs to a single OU process and track the network performance on the corresponding delay task. We then varied both the plasticity timescale and the task delay, while setting the input correlation time proportional to the delay with a factor of 0.5 (task is slower than input), 1 or 2 (task is faster than input). All resulting heatmaps, both for Hebbian (Fig. 3E-G) and anti-Hebbian plasticity (Fig. 3H-J), show a sharp transition from a well-performing to a poorly-performing region. In the tested parameter range, this transition follows a linear relationship closely. We can highlight two main effects for the different ratios. The most notorious one is the progressive degradation of (the optimal) performance for smaller input correlation times *τ*. This is to be expected since the complexity of the task increases. We also note the relatively conserved boundary region for the different ratios. This indicates that the influence of the input correlation timeconstant is small for the determination of the boundary, provided that the plasticity timescale is sufficient to solve the task.

One important point to note is the overall size of range of delays that the network can solve well. For the Hebbian case, the full span of this range is more than an order of magnitude larger than the plasticity timescale *p*. This suggests that a network effect determines the timescale over which information about the input is retained. In a similar way that the recurrence can amplify the timescales of the neurons in the network, the synaptic timescales seem to undergo a similar effect. The resulting outcome is a highly tolerable optimum region that extends way beyond the neuronal and synaptic timescales.

On the other hand, the anti-Hebbian case (*k* = −2) exhibits a similar sharp linear transition, but with a significantly smaller workable region (Fig. 3H-J; see fitted inclination parameters in the Figure caption). It, however, shows slightly better performance in short-delay tasks. This may be a result of the introduction of oscillatory components in the dynamics, as has been shown in Clark and Abbott (2024) and Wakhloo et al. (2025).

### 2.4 Heterogeneous synaptic timescales further optimize performance

Up until now, we have looked at only completely homogeneous networks. In those networks, two intrinsic timescales can be used to process the stimulus: the neuronal timescale, which in our case of memoryless neurons is of the order of the discretization, and the plasticity timescale, which we chose to be more than an order of magnitude larger. We can also ask whether introducing other plasticity timescales could further increase the network performance, given the mixed-timescale nature of our tasks.

There are different ways to introduce heterogeneity to synaptic plasticity. Inspired by our previous results, we choose here to divide the recurrent layer into two groups of neurons, one group projecting slower Hebbian synapses (*p* = 75, *k* = 2), and the other projecting faster anti-Hebbian ones (*p* = 20, *k* = −2). The heterogeneity parameter *γ* gives the proportion of each group, with *γN* slow-projecting and (1 − *γ*)*N* fast-projecting neurons (Fig 4A). At the extremes of *γ*, we get back the two respective homogeneous cases. We also add another, intermediate timescale in the stimulus and task, with the third OU input having correlation scale *τ*_3_ = 60 and the added delay also 60. Therefore, the added input-task timescale also has a corresponding plasticity timescale. What we measure as a result of this setup is a reasonable improvement in performance for intermediate values of *γ*, that is, for networks with some degree of heterogeneity (Fig. 4B-E). When looking at the long-period sample output trajectories, it is difficult to see any meaningful differences between the different cases. However, when zooming in, the differences become more pronounced (Fig. 4C inset). The heterogeneous network is better able to follow the higher-frequency details of the target, whereas the slow homogeneous network does not follow transient deviations on a medium timescale. The same can be seen in the error autocovariance, with the heterogeneous network producing lower values at lower lag (Fig. 4D). Of course, we should expect that the appropriate choices of the synaptic timescales should somewhat follow the characteristics of the input and the task, but even in our simple constructed case, we can see the reflection of those timescales on the performance.

**Figure 4.**
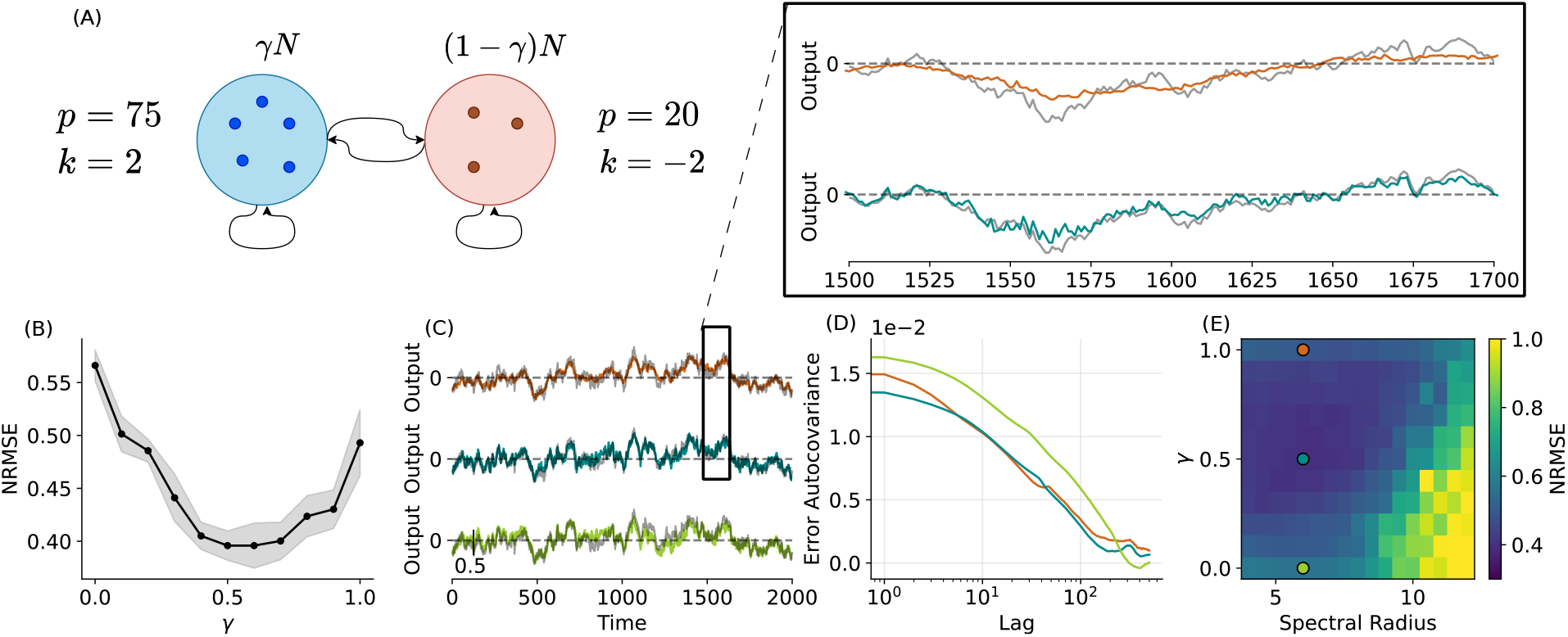
Heterogeneity effects on delay task. (A) Diagram of the designed network. The recurrent layer is divided into two populations with the same connectivity properties. One contains *γN* neurons and projects slow Hebbian synapses, while the remaining neurons project fast anti-Hebbian synapses. (B) NRMSE averaged over 7 realizations of the network and task plotted against the heterogeneity parameter *γ*. (C) Output traces for 3 values of the heterogeneity ratio parameter *γ*. The inset presents a zoomed-in version of the two top traces. (D) Autocovariance of the difference between the output trace and the target. The autocovariance is averaged over all realizations of the network. Note that here we have the autocovariance, not the autocorrelation as in the other figures. (E) NRMSE heatmap for the combinations of *γ* and spectral radius averaged over the realizations of the network and input. Colored dots contain one of the realizations presented in (C). The task consists of the same delay task as in Figure 2A-D, with an added OU input with timescale *τ*_3_ = 60 and a target delay also *d*_3_ = 60.

### 2.5 Number of variables scaling and computational cost

We finally wondered how much of the performance improvement can be attributed to the overall increase in the dimensionality of the dynamical system by adding plastic synapses. For non-plastic reservoirs, adding more neurons to the reservoir usually also improves the resulting task performance, although with diminishing returns (we note, however, that targeted pruning of neurons can improve generalization performance (Dutoit et al., 2009)). We need, therefore, to disentangle the contribution of the size increase and the dynamical contributions.

To this end, we evaluate how the error scales with the number of variables in the system for plastic reservoirs as well as non-plastic reservoirs with Fermi or tanh-neurons. We find that the resulting performance gain is clearly larger even when accounting for the increased size of the system (Fig. 5A-B). For the simpler multi-timescale delay task (Fig. 5A), the performance of a small plastic reservoir (blue curves) is close to that of the largest network of both Fermi (green) and tanh neurons. However, the plastic network continues to improve with size, while the non-plastic ones saturate. We suppose that the initial improvement of the non-plastic networks relates to processing on the fast timescale, but the missing slow timescale keeps the networks from reaching lower error levels (compare Fig. 2M-N). For the more complicated correlated NARMA task (Fig. 5B), the separation between plastic and non-plastic is even greater due to the larger portion of the task dependent on slower processes. Hence, as the synaptic dynamics are the major component responsible for the better memory capacity at slow timescales, increasing the number of neurons does not allow the non-plastic networks to reach the error levels even of the smallest plastic network. We can also note that the performance of the networks with tanh neurons is better in general than that of the non-plastic Fermi neurons for both tested tasks.

**Figure 5.**
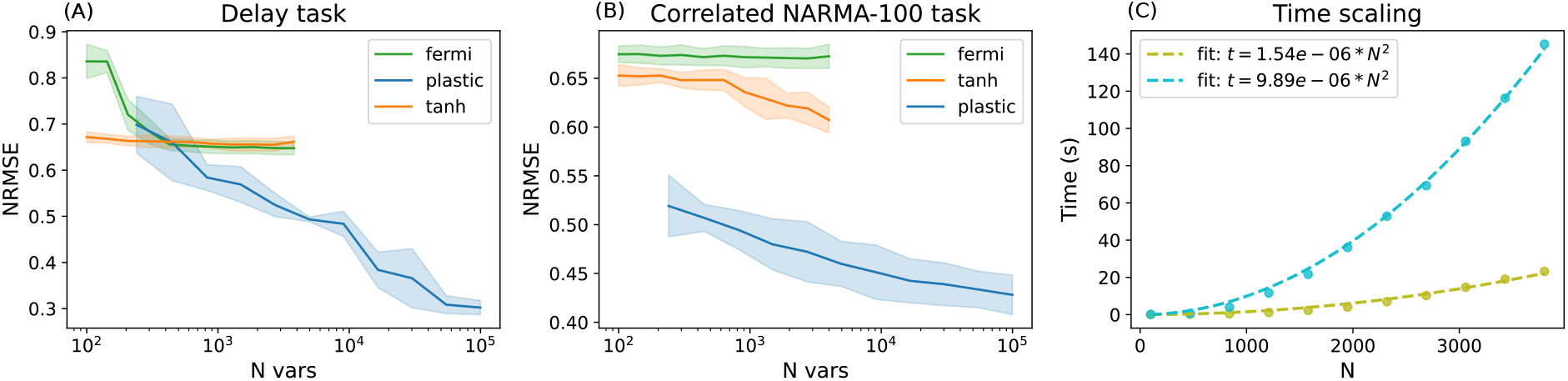
Performance and computational cost scaling. (A)-(B) NRMSE scaling with the number of variables of the models. The models used consist of two fixed-weight networks with Fermi neurons (*ρ*(*c*_*ij*_*J*_*ij*_) = 7) and tanh neurons (*ρ*(*c*_*ij*_*J*_*ij*_) = 0.95), and one plastic network (*k* = 2, *p* = 50, and *ρ*(*c*_*ij*_*J*_*ij*_) = 7). We measured the scaling with the (A) delay task and (B) NARMA-100 task. (C) Time taken to simulate 3000 timesteps of the fixed-weight and plastic network without training (colors as in (A) and (B)). The fitted parabola indicates that computational cost is still of order 2 for both models (*R*^2^ *>* 0.9). Parameters for (A) are as in Fig. 2A-D. Parameters for (B) are as in Fig. 2O-R.

Apart from the number of dynamic variables, also the scaling of the computational costs with the network size *N* is important for more complex applications. Strikingly, it can be expected that the additional cost of adding plasticity for running the reservoir is only of order *O*(*N* ^2^), since there is only an additional outer product to compute in the term *ϕ*_*i*_(*t*)*ϕ*_*j*_(*t*). However, the computational cost of the static reservoir is already of order *O*(*N* ^2^), so the plasticity doesn’t increase the complexity order of the algorithm. We tested this by running the plastic and non-plastic reservoirs of different sizes with a given input and calculating the computing time required. Fitting a parabola to both datasets clearly indicates the quadratic scaling, with the increase in computational cost coming only as the quadratic factor (Fig. 5C; *R*^2^ *>* 0.9). We would like to point out that we are here disregarding the higher computational cost of training networks with more neurons that involve matrix inversions.

### 2.6 Conditions for observing plasticity-induced performance gains

We showed that the introduction of the plastic Hebbian term improved the performance of our reservoir on the tested memory tasks. We next wanted to identify which components of our model are needed for the improvements to emerge. For the plasticity rule, this poses the question of whether they emerge from the presynaptic dependence, the postsynaptic dependence, or the cooperation of both terms in the Hebbian rule. To this end, we first apply the same delay task protocol as in Figure 2A-D, using a Hebbian rule, a postsynaptic-only dependent rule (Post-plastic), and a presynaptic-only dependent rule (Pre-plastic) (see Table 1). We find that the performance is similar for both the Hebbian and the Post-plastic case, but for the Pre-plastic model it is mostly degraded to the non-plastic levels (Fig. 6A-C). Although some marginal effects of the plasticity are still present in the Pre-plastic model, we therefore hypothesized that a presynaptic-only dependent plasticity model is not able to carry the slower temporal information in a manner that is readable by an output layer, and the relevant dependence is the postsynaptic one.

**Table 1:** Plasticity models used and the differences between them. *C*(*t, s*) is here the autocovariance of the neurons and *µ*(*t*) its mean.

| Model | Update rule | Kernel | Post-synaptic term |
| --- | --- | --- | --- |
| Hebbian | $A_{ij}(t+1) = (1 - \frac{1}{p})A_{ij}(t) + \frac{k}{Ncp}\phi_i(t)\phi_j(t)$ | $K(t, s) = e^{-\frac{t-s}{p}}C(t, s)$ | $\psi(t) = \phi(x(t))$ |
| Post-plastic | $A_{ij}(t+1) = (1 - \frac{1}{p})A_{ij}(t) + \frac{k}{Ncp}\phi_i(t)$ | $K(t, s) = e^{-\frac{t-s}{p}}\mu(t)$ | $\psi(t) = \phi(x(t))$ |
| Pre-plastic | $A_{ij}(t+1) = (1 - \frac{1}{p})A_{ij}(t) + \frac{k}{Ncp}\phi_j(t)$ | $K(t, s) = e^{-\frac{t-s}{p}}C(t, s)$ | $\psi(t) = 1$ |

**Figure 6.**
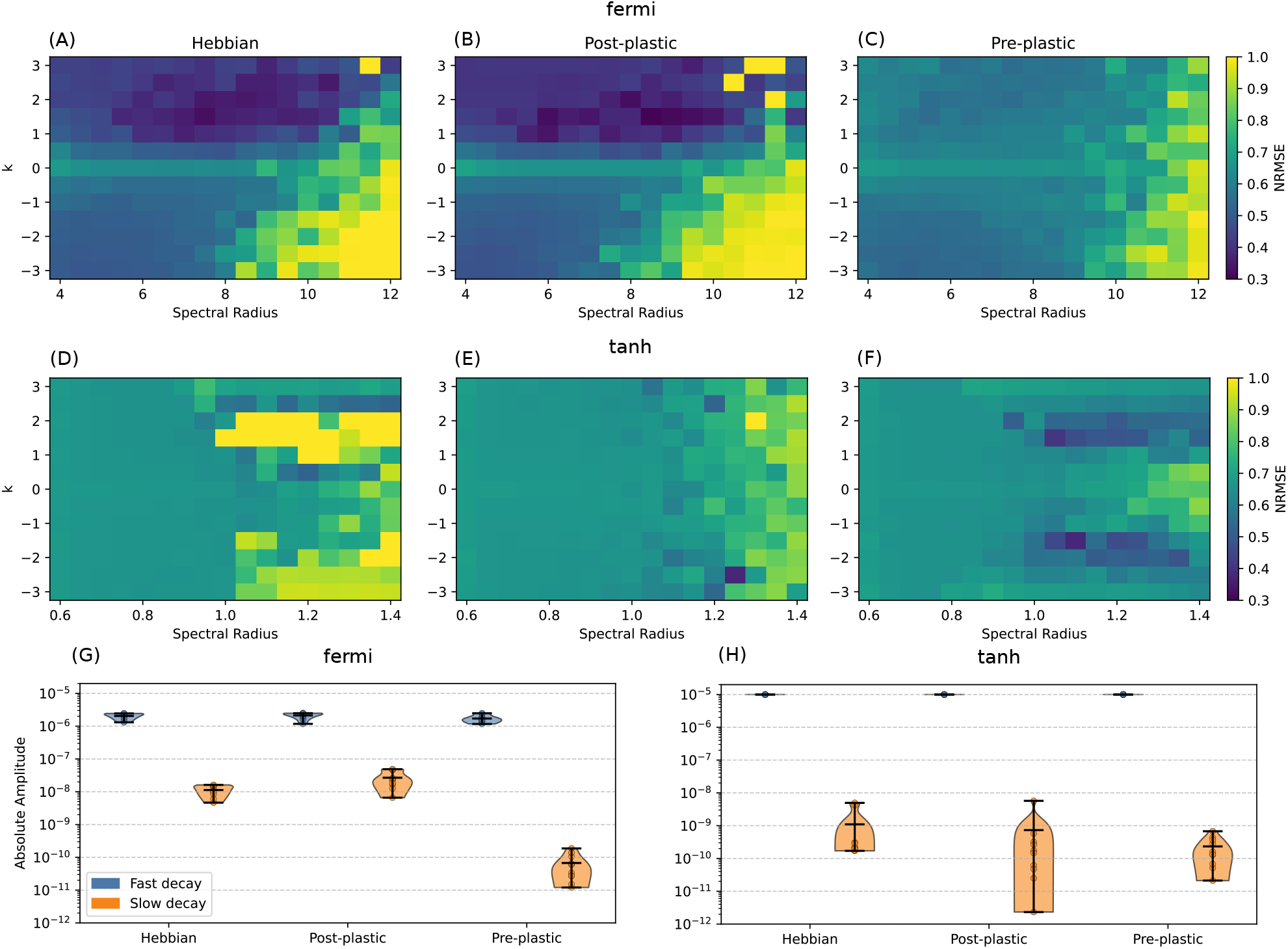
Performance and timescale comparison for different plasticity models. (A)-(C) NRMSE heatmaps for the (A) Hebbian plasticity model, (B) postsynaptic-dependent plasticity, and (C) presynaptic-dependent plasticity using Fermi neurons. (D)-(H) NRMSE heatmaps for the (A) Hebbian plasticity model, (B) postsynaptic-dependent plasticity, and (C) presynaptic-dependent plasticity using hyperbolic tangent neurons. Parameters as in Fig 2. (G)-(H) Fitting amplitudes using a double exponential function for the difference between a locally perturbed single neuron trajectory and an unperturbed one. We used a fully connected network (without self-connections), time step of 10^*−*3^ and a perturbation input of 10^*−*4^, using 10000 timesteps for the fitting. (G) implements a network of fermi neurons with spectral radius *ρ*(*J*_*ij*_) = 1 and (H) implements a hyperbolic tangent network with *ρ*(*J*_*ij*_) = 0.1

Following this hypothesis, we investigated the transients our model shows in response to small perturbation currents. To do this, we first consider the continuous, autonomous, fully-connected version of our system:

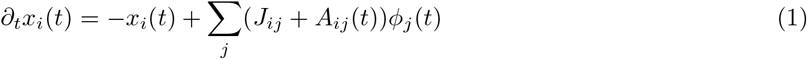

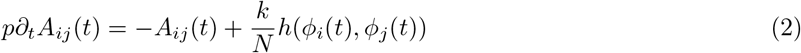

where *h*(*ϕ*_*i*_(*t*), *ϕ*_*j*_(*t*)) indicates one of the three models above, that is: *h*(*ϕ*_*i*_(*t*), *ϕ*_*j*_(*t*)) = *ϕ*_*i*_(*t*)*ϕ*_*j*_(*t*) for the Hebbian one; *h*(*ϕ*_*i*_(*t*), *ϕ*_*j*_(*t*)) = *ϕ*_*i*_(*t*) for the Post-plastic; and *h*(*ϕ*_*i*_(*t*), *ϕ*_*j*_(*t*)) = *ϕ*_*j*_(*t*) for the Pre-plastic. Considering that the system has reached equilibrium and is experiencing a small perturbation on a single neuron at time *t*_0_, we can use dynamic mean field theory (DMFT) to arrive at a treatable, one-dimensional model description (Clark and Abbott, 2024) and derive the following equation for the linear response of the system (see Methods for a detailed derivation):

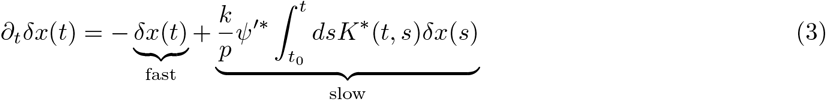

where *δx*(*t*) is the deviation from the fixed point *x*^*∗*^, *K*^*∗*^(*t, s*) is the kernel term evaluated at the fixed point and depends on the model (see Table 1), and *ψ*^*′∗*^ is the derivative of the postsynaptic term with respect to *x* at the fixed point (also dependent on the model). We can see from this equation that two terms contribute to the response of the network, one fast term and one slow (given large *p*). Since the Pre-plastic model does not depend on *ϕ*(*t*) (*ψ*(*t*) = 1), the derivative *ψ*^*′∗*^ in the slow response term is zero, such that the timescale extension effects of the plasticity vanishes. Note, however, that the above equations were derived for infinite-sized networks. For finite-size networks, the response term exhibits a slow timescale correction (given the implicit dependence of the kernel *K* on *x*) without *ψ*^*′∗*^, explaining the marginal performance gains of Figure 6C (see also Methods). However, we conclude that our perturbation analysis supports the hypothesis that the improvement for slow timescales emerges from a postsynaptic dependence.

When examining the above equation, we note that for *p*≫ 1 and *p*≫ *k*, the network exhibits essentially two decay rates. For shorter *t*, the integral is small and since *k/p <<* 1, the fast decay term dominates, leading to a decay with timescale *τ*_fast_ = 1. For longer *t*, the integral term generates finite contributions that start to dominate the fast decay. Based on this reasoning, we can test our hypothesis also in simulations using a fully connected network (without self-connections) with spectral radius *ρ*(*J*_*ij*_) = 1, in order to minimize nonlinear and finite-size effects. In particular, we simulated each network twice, introduced a small perturbation at one neuron at a time *t*_0_, and then tracked the difference in the neural activities between simulations. We then fit the following functions to that difference:

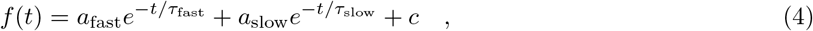

where the faster decay term *τ*_fast_ corresponds to the neuronal leak and should be equal to 1 for all models, and the slower decay coefficient *τ*_slow_ is the result of the activity response term. The observed decay time constants match our theoretical expectations, with the neuronal decay averaging 1 and similar amplitudes for all the cases (Fig. 6G). This fast decay explains why the network is still able to appropriately account for the short delay in the mixed-timescale task. The typical neuronal timescales present in the non-plastic networks, as shown by Crisanti and Sompolinsky (2018), are still accessible to the output layer in the plastic ones.

The effect of the postsynaptic term in the plasticity rules is noticeable in the slower decay constant for the single-neuron perturbation *τ*_slow_. For all cases, the observed value is of order 10^2^; however, the amplitudes for the Hebbian and Post-plastic cases are two orders of magnitude larger than the Pre-plastic network. Hence, the slower timescales generated by the plasticity dynamics are much less noticeable in the Pre-plastic network, as they come mainly through finite-size effects (autocovariance response). Overall, this gives us an explanation for why both the Hebbian and the Post-plastic plasticity models exhibit similar improvements for the mixed task, and the Pre-plastic only sees marginal improvements.

The form of the perturbation dynamics in equation (3) also reveals a condition for the neuronal dynamics. If we choose a transfer function *ϕ*(*x*(*t*)) that gives a stable fixed point at zero (as is the case of the widely used *ϕ*(*x*(*t*)) = tanh(*x*(*t*))), this zero fixed point concentrates the activity distribution into a delta function at zero, making both the mean *µ*(*t*) and autocovariance *C*(*t, s*) zero, and consequently *K*^*∗*^(*t, s*) as well. This is not the case for non-zero fixed points, which makes their distribution heterogeneous in general. Therefore, the perturbation is nullified by the kernel term and the slow timescale should not be present. We tested this by repeating our simulations for the tanh(*x*) transfer function shown in Figure 6D-F. As predicted, the plasticity shows basically no effect in the stable region, only exhibiting different behavior in the chaotic regime where the fixed point at zero becomes unstable, and the kernel is no longer null. Furthermore, when looking at the perturbation decay responses, we can observe very small slow amplitudes comparable to the Pre-plastic case for the Fermi neurons, while the fast amplitude is a little bit higher (Fig. 6H). This indicates the dominance of the neuronal timescales over the plastic ones, making the slow part not readable by the output layer.

In summary, we provided a theoretical understanding and simulation evidence that a postsynaptic dependence of the plasticity and a neuronal transfer function leading to a non-zero activity fixed-point are the critical components to achieve plasticity induced performance gains on multi-timescale time-series processing.

## 3 Discussion

We have demonstrated that plasticity mechanisms improves reservoir computing for tasks with mixed-timescales. In particular, we showed that plasticity extends the network’s memory range while maintaining its short-timescale capabilities. This memory extension is based on weight dynamics as a fundamental resource rather than from the convergence of the weights to an optimal fixed-point, as freezing the recurrent weights and retraining the output layer resets the performance to baseline non-plastic levels. We can understand this lack of change in the frozen-retrained network by looking at the scaling of the static weight term *J* and the plastic weights *A*. As argued in Clark and Abbott (2024), the magnitude of *J* scales with 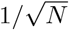 while *A* scales as 1*/N*. Changes in the network’s weight structure are therefore only marginal, vanishing as its size increases. The learning rule can’t meaningfully explore the weight space to create clusters of strongly connected neurons as in typical cell assemblies (Zenke et al., 2015). The relevant contribution comes only after summing over all the incoming synapses in the dynamical regime, where the effects appear as a filtered correlation function. Hence, the scaling properties of the rule provides only meaningful dynamical effects.

In the biological literature, plasticity, especially of the Hebbian type, is typically conceived as a fixed-point optimization process for synaptic weights; that is, the weights converge to a specific configuration that optimizes the proposed computation (Hopfield, 1982; Zenke et al., 2015). This view arises primarily in the context of cell assemblies, where learning is framed within an attractor paradigm in which stable patterns of activity are encoded in the recurrent weights of the network. Our work provides a complementary view of synaptic plasticity, where the timescales of the plasticity can be used as computational resources to be taken advantage of. These timescales can serve as a bridge between the typical fast neuronal dynamics, of the order of milliseconds, and the behavioral range. Experimental work suggests a possible role of plasticity in keeping information in working memory (Wolff et al., 2017), while some models propose mechanistic explanations using short-term plasticity (STP; cf. Masse et al. (2019); Mongillo and Tsodyks (2026)). One recent example uses the longer timescales of synaptic augmentation together with standard STP to encode sequence information (Mongillo and Tsodyks, 2026). Taken together, this implies that the here presented principles may not only be biologically inspired, but also employed by the brain to solve multi-timescale task. When looking at our model parameters, namely the spectral radius, plasticity amplitude, and plasticity timescale, we have three control knobs that tune different components of the network: The main effect of the *spectral radius* is to define the transition from steady state to chaos, as originally described in Sompolinsky et al. (1988), which results in a drop in task performance. Naturally, the exact transition boundary also depends on the input (Rajan et al., 2010) and on the plasticity strength (Clark and Abbott, 2024). This transition is also related to the standard prescriptions for echo state networks to exhibit the echo state property, requiring stable dynamics (Jaeger, 2001; Lukoševičius and Jaeger, 2009). A secondary effect is the performance improvement in the Hebbian (*k >* 0) networks for the mixed delay task (Fig. 2D). We speculate that this improvement comes from the spread of the network neuronal timescales (Manneschi et al., 2021), which enable the fast-timescale computations in our tasks, which are up to an order of magnitude slower than the intrinsic neuronal timescale. This is also consistent with earlier findings that criticality improves performance on some tasks (Cramer et al., 2020; Boedecker et al., 2012; Legenstein and Maass, 2007).

On the other hand, the *plasticity amplitude k* has a more distinct effect on the network performance. It indicates how much temporal correlation and past states are injected into the neural dynamics as a current, whether positive or negative. The task error initially rapidly decreases with *k* to an optimum and then saturates at a close value. There is, however, a difference in the optimum obtained for positive and negative synaptic strength (Hebbian and anti-Hebbian). This is mostly related to the timescale range of each plasticity type. As we have shown, anti-Hebbian plasticity behaves better when processing shorter timescales, exhibiting a more restricted range. Meanwhile, the Hebbian case displays a wider well-performing region that encapsulates a wide range of middle to slow timescales (Fig. 3).

When looking at the *plasticity timescale p* parameter, our network exhibits a wide optimum range for the tested task. This indicates that the choice of the timescale parameter is robust to changes in the input and task configuration. This is even clearer for the single-input case, where the optimum range extends more than an order of magnitude beyond the value of the plasticity timescale (Fig. 3). We note that, explicitly, the model has only two timescales in it: the neuronal one on order one timestep (implicit in the choice of leak), and the plastic one *p*. With just those two choices, we were able to significantly decrease the error in both tested tasks involving mixing of timescales, including the correlated NARMA-100 that nonlinearly requires information from all previous 100 timesteps. However, a deeper investigation of the parameter tuning in more complex settings is also required, since it is already known that tuning the leak rate can be challenging (Jaeger et al., 2007).

We also observed that heterogeneous plasticity timescales in the form of mixed populations of neurons with different synaptic properties can optimize the performance of the network by adding more timescales. By mixing a population with slow Hebbian plasticity and a fast anti-Hebbian plasticity, we were able to construct a network better capable of capturing the imposed three timescales of the task. It is important to note that we used the simplest form of heterogeneity here. Given the large optimum region for the homogeneous cases and the timescales used in the task, the addition of more populations had progressively less impact on performance (data not shown). Thus, biological neuronal networks may only need a few plasticity mechanisms at different timescales to serve a broad range of multi-timescale tasks (Tetzlaff et al., 2012). Heterogeneity in the neuronal timescales has been shown to improve reservoir processing capabilities in tasks involving fast and slow chaotic systems (Tanaka et al., 2022), generate better and more stable results under training of the recurrent weights Perez-Nieves et al. (2021), and provide networks better able to handle a diverse set of tasks (Golmohammadi et al., 2025). It is an interesting question whether those properties also translate to heterogeneity in the plasticity timescales for tasks on longer or multiple timescales.

Importantly, we notice that the performance gains for the proposed memory tasks come mostly from the postsynaptic dependency of the plasticity rule. This dependency appears as a memory term filtered by an exponential kernel also dependent on the autocovariance. As a result, the system exhibits a second decay timescale besides the neuronal decay rate, which is also present in the network dynamics (Fig. 6). The mixing of timescales in the task is then properly handled by the two timescales in the network: the one generated by the neuronal dynamics, and the one generated by the filtered postsynaptic plasticity dependence. In this sense, a plasticity rule that is postsynaptic-only dependent also has similar performance to the Hebbian one. When using postsynaptic-only dependent synaptic plasticity, we can also expect a change in the computation cost compared to Hebbian plasticity. The postsynaptic-dependent plasticity does not involve the outer product computation that adds the order *O*(*N* ^2^) cost but is purely linear. Thus, adding postsynaptic-only dependent plasticity adds just a sub-leading term to the scaling of computational cost with network size, while retaining the same improvements for reservoir computing.

Of course, this doesn’t mean that Hebbian plasticity plays no role in processing timeseries beyond the reservoir framework. For example, as shown by Wakhloo et al. (2025), the Hebbian structure can create and sustain persistent oscillations with the same characteristics of the oscillating transient input injected into the network. This behavior emerges from the interaction of the input, the static recurrent weights, and the plastic ones. Thus, we would expect to observe a more pronounced performance increase for networks with Hebbian plasticity in tasks with oscillatory components. Also, for other kinds of tasks, it may be possible that Hebbian plasticity can take advantage of a non-random composition of the static weights *J*_*ij*_ in a similar fashion in other types of tasks.

Furthermore, since we here focused on the timescale properties of plasticity, we limited ourselves only to small variations of the Hebbian formulation. In reality, there are a plethora of different learning rules proposed in neuroscience and machine learning, with different properties and tackling different problems (Gerstner and Kistler, 2002; Hennig, 2013; Tetzlaff et al., 2011; Toyoizumi et al., 2005). We expect that the improvements observed here may be present for the rules with postsynaptic dependence and networks with the nonzero fixed-point property. However, the exact properties of the temporal processing may change and may give rise to even more complex computations on the slow timescale. For example, a nonlinear dependence of the weight change on the postsynaptic activity will change the kernel term, and therefore the filtering property of the neuronal activity. Similarly, even more complex synaptic plasticity mechanisms such as cascade models (Fusi et al., 2005; Benna and Fusi, 2016) can introduce other slower processes that will be integrated into the kernel. The exploration of those properties can be fruitful for extracting different temporal attributes of a given stimulus.

Another necessity for the here-demonstrated performance improvements is the nonzero heterogeneous fixed-point of the neural activities, which makes the temporal dynamics of the plastic weights readable. When the network exhibits a homogeneous fixed-point at zero, the kernel term that appears as a factor in the postsynaptic response term is also zero, erasing the effect of the perturbation. This is the case of the standard reservoir formulation with tanh neurons (Jaeger, 2001; Tanaka et al., 2022; Ozturk et al., 2007; Wakhloo et al., 2025). As we have shown, for tanh neurons, the plasticity has basically no effect on the network performance, making local perturbations unnoticeable to the output layer. Hence, to observe the same performance improvements with tanh neurons, adding heterogeneity to the currents in the form of a bias for each neuron, or making the network unbalanced by changing the mean of the static connectivity, would be an option worth exploring.

Overall, we have shown that the dynamical properties of synaptic plasticity can serve as a computational resource for processing stimuli across multiple timescales. By combining fast neuronal dynamics with slower synaptic dynamics, the network preserves its intrinsic neuronal timescales while gaining access to additional temporal structure through plasticity. This provides a simple and computationally efficient mechanism for multi-timescale processing, with potential applications in neuromorphic signal processing. More broadly, our results suggest that synaptic plasticity may play a computational role beyond adapting network connectivity, acting instead as a dynamical element that extends the temporal range over which biological neural circuits can process and retain information.

## 4 Methods

### 4.1 Network model

We consider an echo-state network architecture consisting of an input, a recurrent, and an output layer. Inputs consist of *N*_*I*_ Ornstein-Uhlenbeck (OU) processes with statistics

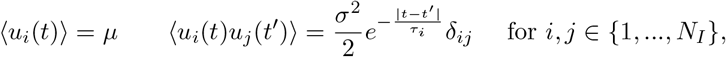

where the angle bracket ⟨…⟩ represents an average over the distribution of the process. In all the cases used in this work, we choose *µ* = 0 and *σ* = 0.5. Correlation timescales *τ*_*i*_ are specified differently for each of the tasks performed (see Figures).

The recurrent layer is composed of *N* neurons sparsely connected, obeying the following dynamics:

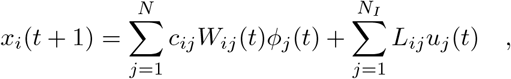

with *c*_*ij*_ drawn from a Bernoulli distribution with probability *c* = 0.1, *L*_*ij*_ drawn from an uniform distribution in the interval (−0.5, 0.5), and the nonlinear transfer-function given by a fermi function

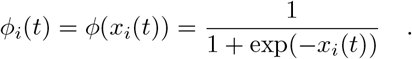

In Figure 5, we also used a hyperbolic tangent for comparison. Note that the dynamics don’t contain a leak term, so that in the context of an Euler discretization with timestep *dt* of the equivalent differential equation, we would have *dt* = *τ*_*L*_. We will consider the time *t* in units of *τ*_*L*_ (that is, choosing *τ*_*L*_ = 1), such that *dt* is constrained to 1. Therefore, we also used *dt* = 1 for the generation of the OU process. The recurrent weights follow similar dynamics as described by Clark and Abbott (2024). It is composed of a static part *J*_*ij*_ drawn from a Gaussian distribution with zero mean and fixed spectral radius *ρ*(*c*_*ij*_*J*_*ij*_), and a plastic part *A*_*ij*_. More specifically:

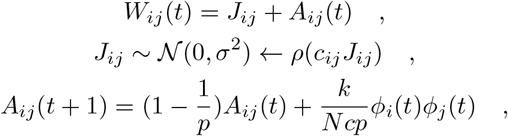

where *k* is the plasticity strength, *p* is the plasticity timescale, and *c* the sparsity as defined previously. Note that, since *t* is in units of the leak rate *τ*_*L*_, *p* also has the same units meaning that *p* is measured in multiples of *τ*_*L*_.

### 4.2 Tasks

Two tasks were used to evaluate the network capacity to process mixed-timescale inputs: a multiple timescale delay task and a correlated NARMA-100 task. The delay task consists of one target output constructed as the average of delayed versions of the inputs, that is,

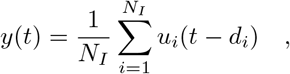

where *d*_*i*_ is the delay on input *u*_*i*_(*t*). Three setups were considered for this task: The first one used two inputs, one fast (*τ*_1_ = 20) and one slow (*τ*_2_ = 400), with respective delays *d*_1_ = 20 and *d*_2_ = 400. In this sense, we asked the network to remember a more distant past from the slower stimulus than the faster one. We also added a third timescale for the heterogeneous task in order to make the heterogeneous effects clearer, with *τ*_3_ = 60 and *d*_3_ = 60. In the third setup, where we evaluated the overall memory range of the network given the synaptic timescale *p*, one single input was used, keeping the ratio of delay and correlation constant at either 0.5, 1 or 2.

The correlated NARMA-100 task considers nonlinear products over its past values. The specific form used in this work is given by

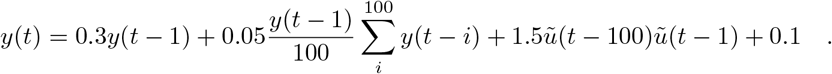

Usually, the input for a typical NARMA task is uncorrelated noise. However, since we are interested in testing the ability of the network to use temporal correlation in the input to perform mixed-timescale computations, we used a different setup. The network received four separate OU inputs with correlations: *τ*_1_ = 10, *τ*_2_ = 30, *τ*_3_ = 70, and *τ*_4_ = 100. The input to the target *ũ*(*t*) is then constructed as

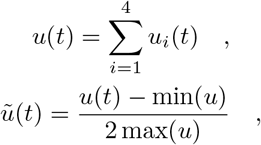

with min(*u*) and max(*u*) being the minimum and maximum value of the sum *u* over the whole duration, of the simulation respectively. This normalization between 0 and 1*/*2 is done for stability reasons.

### 4.3 Training and testing

For training and testing procedures, we employed the normalized root mean squared error (NRMSE) (Lukoševičius and Jaeger, 2009). Since our outputs consist of single neurons, the resulting formula is:

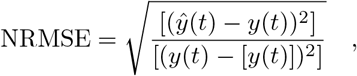

where *y*(*t*) is the target output, *ŷ*(*t*) is the output estimated by the network, and the brackets […] represent the average over time. The estimated output is the result of a simple linear operation:

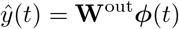

where **W**^out^ is the 1×*N* matrix that we want to train. Training is performed by a Tikhonov/Ridge regression:

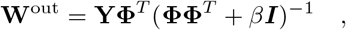

with **Y** being the 1 × *T* matrix of concatenated times [*y*(*t*_0_), *y*(*t*_0_ + 1),…,*y*(*t*_0_ + *T* − 1)], **Φ** is the *N* × *T* concatenated state-vector matrix [***ϕ***(*t*_0_), ***ϕ***(*t*_0_ + 1),…,***ϕ***(*t*_0_ + *T* −1)], *I* is an *N* ×*N* identity matrix, and *β* is the regularization parameter (here chosen to be *β* = 10^*−*7^).

The standard training and testing protocol is done in the following way: we first present 20000 timesteps of the input to the network for it to equilibrate. We follow up by presenting the next 100000 timesteps and use corresponding state vectors ***ϕ***(*t*) generated by the input to train the output weights **W**^out^. Testing is then done by calculating the error over another 20000 timesteps of the input. Heatmaps are constructed by averaging the NRMSE of different realizations of the network and inputs.

### 4.4 Frozen weights protocol

To evaluate whether the performance improvements result from plasticity dynamics or fixed-point optimizations, we used a frozen-weights protocol (compare Clark and Abbott, 2024). After a standard round of equilibration, training, and testing, the plastic synaptic terms are fixed at their last updated value, and another round of testing is performed. We then calculate the delta error from the frozen period and the previous plastic testing phase. This delta is averaged over three realizations of the network and input. After testing on the static weights, we retrain the output layer over a period of 100000 timesteps and re-test again for 20000.

### 4.5 Perturbation derivations

Our aim here is to derive a formal equation for the response of the system to a small perturbation. We do note that using an appropriate numerical scheme for solving the resulting set of self-consistent equations as done by Clark and Abbott (2024), theoretical values for the decay rates are obtainable.

We start our derivation with the continuous, autonomous, fully connected model:

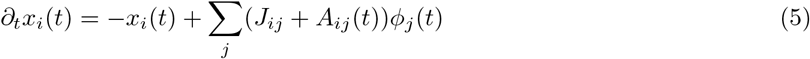

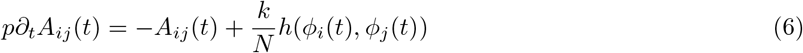

with

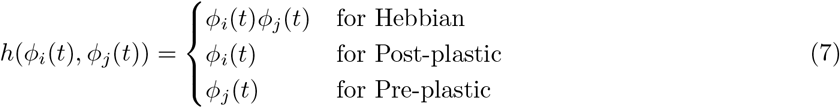

Integrating *A*_*ij*_ results in

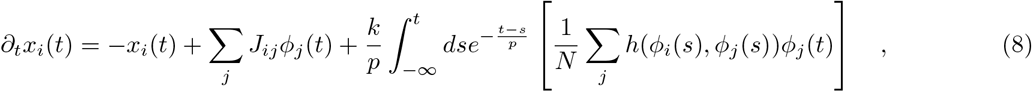

We can already note that the term in brackets will produce an averaging quantity, being it the average activity (in the Post-plastic case) or the autocovariance (in the Hebbian and Pre-plastic cases). This allows us to write the equivalent DMFT system Clark and Abbott (2024):

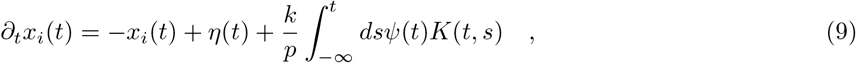

where *ψ*(*t*) and *K*(*t, s*) are defined as in Table 1. Since the relevant phenomena occur in the stable regime in our case, we can look at what happens around the fixed-point solution *x*^*∗*^. We consider the single-site linear response properties of the network, that is, we look at the response of a single neuron given a small perturbation only on this neuron. Performing a perturbation in a single neuron around the fixed point at time *t*_0_, and linearizing the dependent quantities

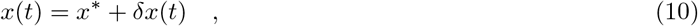

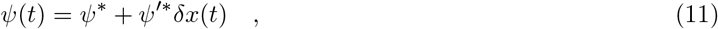

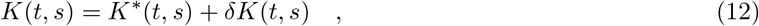

where stars represent fixed point values. Substituting those expressions into (9) and keeping only linear terms, we can finally obtain the differential equation for the linear response of the system

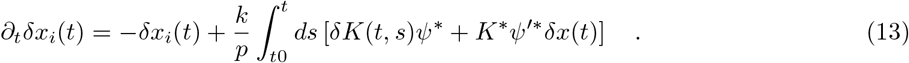

In the large *N* limit and giving the single site nature of the perturbation, the contribution *δK*(*t, s*) goes to zero, since it is an averaging quantity and exhibits an 1*/N* dependency. This is to be expected, since in the DMFT description this quantity is treated independently of the values of *x*(*t*). We do show this step, however, to point out that for finite networks, there is still some contribution from the perturbation in the kernel term. Not only that, but for global perturbations, the kernel response becomes finite in the mean-field limit. We also note that a global perturbation would result in a finite contribution of the kernel term. Thus, taking this limit for our local perturbation equation, we get the expression presented in the results section:

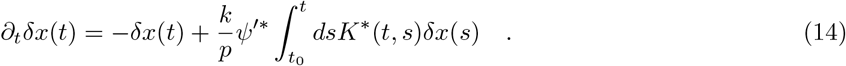

Note, we can split the 1D linear response equation into a two-dimensional system where we have again a plasticity related variable. By noting that the fixed-point kernel term is just an exponential decay times a constant, we can finally write

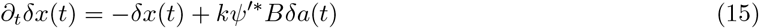

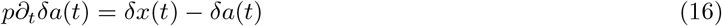

where *B* is the fixed point value of the average or the autocovariance of the activity (given that we are looking at perturbations around the stationary fixed-point), depending on the model. The resulting decay dynamics are explicitly 2D, with the main contribution to the plasticity variable coming from the neuron activity. This also provides another way to look at the double exponential decay behavior that arises when *p* is sufficiently large. In this regime, we can use separation of timescales, so we can first treat *a* as constant and get an exponential decay for *x* and then solve separately for *a* to get another exponential decay. Both contribute independently to the two decay timescales.

### 4.6 Perturbation simulations and fitting

To estimate the decay rates of the different plasticity models, we used a small perturbation protocol and fitting. To this end, we used networks composed of 1000 fully connected neurons (except for self-connections *J*_*ii*_ = 0), setting the spectral radius *ρ*(*J*_*ij*_) = 1 for the Fermi neurons network and *ρ*(*J*_*ij*_) = 0.1 for the tanh network (that is, we used 10% of the observed critical value 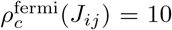 and 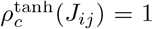 in order to minimize nonlinear and finite-size effects in the resulting traces). The network receives no input besides the perturbation over the whole simulation period. To get stable traces, we consider the continuous version of the system mentioned above (Equations 5 and 6). For the simulations, we used Euler discretization with *dt* = 0.001. We ran the network over 100000 timesteps to equilibrate, copied the state of the network, and ran a perturbed and an unperturbed version of the same network for 10000 timesteps. The perturbation is done by adding a small current with amplitude *ϵ* = 10^*−*4^ to a single neuron. The trace to be fitted is then the difference between the perturbed and unperturbed single neuron trajectory.

For the fitting part, we consider the double exponential function:

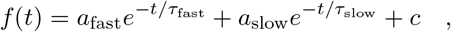

where *a*_fast_ and *τ*_fast_ belong to the fast-decaying part corresponding to the neuronal timescale, *a*_slow_ and *τ*_slow_ correspond to the slow synaptic response, and *c* is a constant. The fittings were only accepted if the *R*^2^ measure was greater than 0.99, that is:

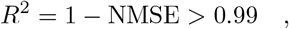

with the NMSE calculated between the fitted function and the resulting simulated trace as

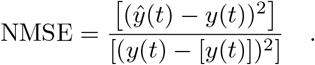

